# “Lethal effects of the wild potato *Solanum bulbocastanum* on the planthopper *Pentastiridius leporinus*, a vector of bacterial pathogens in potato”

**DOI:** 10.1101/2025.10.12.681902

**Authors:** Eva Therhaag, Karin Gorzolka, Jürgen Gross

## Abstract

The planthopper *Pentastiridius leporinus* [Hemiptera: Cixiidae] is a vector of the two plant pathogenic bacteria ‘*Candidatus* Arsenophonus phytopathogenicus’ and ‘*Candidatus* Phytoplasma solani’ in potato. An infection leads to symptoms such as rubbery and non-processable tubers and consequently to substantial decreases in yield and tuber quality as well as to abandonment of seed potato production in affected areas. Up to date, resistant potato varieties are not yet known, and the tool box of integrated pest management to prevent the spread of *P. leporinus* as causal vector is quite limited. Mortality experiments designed as no-choice trials on potato cultivar ‘Gala’ (*Solanum tuberosum*) and on its wild relative *S. bulbocastanum* revealed significant effects of the latter on the survival rate of the vector. LC/MS analysis of the insects and their intestines showed that acyl solamines were retrieved only in individuals from the *S. bulbocastanum* plants. Furthermore, choice-tests were carried out to study the vector behavior. A preference towards *S. bulbocastanum* vs. *S. tuberosum* ‘Gala’ was observed despite its lethal effects. Volatile organic compounds of the two different species were collected and analyzed by GC/MS. The two profiles differed in 39 of 80 compounds. The wild potato species *S. bulbocastanum* and its compounds are discussed as potential starting points for further research on sustainable management of *P. leporinus*.

**Key Message:**

- We tested a wild potato against a planthopper which vectors pathogenic bacteria to potato
- Wild potato led to higher mortality of the planthopper but was preferred against cultivated potato
- Uptake assays suggest that the phloem sucking planthopper is taking up xylem sap as well
- Acyl solamines were only found in guts from planthoppers sucking on the wild potato
- Our results present a starting point for new approaches for sustainable vector control

## 1 Introduction

The history of potato (*Solanum tuberosum* L.) cultivation dates back 7,000 years. Potato belongs to the first domesticated food crop and still plays an important role in human diets and nutrition – it is grown on each continent despite Antarctica (Zaheer and Akhtar 2016; Bakhsh et al. 2023). In 2023, the global potato production was estimated about 383 million tons (FAO 2024). A bit more than a twelfth of that, 48.3 million tons of potatoes were harvested in Europe. There, the five countries Germany, France, the Netherlands, Poland and Belgium made up almost 70% of the harvested area (Eurostat 2024).

Since 2022, the potato production in the Southwest of Germany encounters severe challenges. In particular, in Hesse, the production of seed potato decreased dramatically over the last few years from 290 ha in 2021 to 67 ha in 2024 (LLH 2024). This coincidences with the first finding of a relatively new vector in this crop, the planthopper *Pentastiridius leporinus* L. [Hemiptera: Cixiidae] (Behrmann et al. 2023). It transmits the bacterial pathogens ‘*Candidatus* Phytoplasma solani’ and ‘*Candidatus* Arsenophonus phytopathogenicus’ to potato (Behrmann et al. 2023; Therhaag et al. 2024a). These transmitted pathogens have been reported recently as an emerging risk in a stakeholder discussion meeting of the European Food Safety Agency (EFSA 2024).

An infestation of the crop may be associated with symptoms of wilting, yellow to purplish leaf tops, rubbery, small or airy tubers and loss of yield (Therhaag et al. 2024a; Rinklef et al. 2024). In tubers, infection with the bacteria leads to crop losses, partly due to reduced storage, processing or germination capacity (Lindner et al. 2011; Mahillon et al. 2025; Lang et al. 2025). Since adult *P. leporinus* have only been reported quite recently as vector of the afore mentioned pathogens in potato (Therhaag et al. 2024a), specific vector management options in potatoes have not yet been studied. In 2024, other crops and vegetables have been reported to be infested by one or both of the transmitted pathogens (Therhaag et al. 2024b; Lang et al. 2025), which aggravates the situation in areas of high abundance of *P. leporinus*. This underlines the urgent need to investigate and develop short- and long-term management tools. Since the planthopper is well adapted to the cropping rotation, which includes winter cereals sown after potato or sugar beet, a starvation period due to fallow, maize or other summer crops may be a promising attempt to break its lifecycle and reduce the population (Bressan et al. 2010; Pfitzer et al. 2024). An intercrop that has a negative impact on the planthopper could also be included in the crop rotation.

Besides the lever of an adapted crop rotation, another important part of integrated pest management (IPM) is the use of resistant or robust varieties, in particular when bacterial plant pathogens are transmitted by polyphagous vectors across different crops (Agrios 2005). In the case of potato, various wild relatives are available which could serve as a source of desirable traits such as resistance or resilience to better cope with pests and diseases (Jansky and Spooner 2018). Efforts have been made to introduce resistance genes from *Solanum* species to domesticated potato as for example reported for potato late blight caused by *Phytophthora infestans* (Malcolmson and Black 1966; Van Der Vossen et al. 2003; Gopal 2023) or the Colorado potato beetle (*Leptinotarsa decemlineata*) (Crossley et al. 2018). However, no resistance traits against *P. leporinus* or the transmitted pathogenic bacteria were found in cultivated or wild potato species until today.

Recently, bioassays with a bioactive compound isolated from the wild relative *Solanum bulbocastanum* Dun. showed effects against the Colorado potato beetle (*L. decemlineata*), the potato flea beetle (*Epitrix papa*), and *P. infestans* (Gorzolka et al. 2025). The isolated compound (C16:3(7Z,10Z,13Z)-solamine) is an acyl polyamine. An acyl solamine fraction obtained from *S. bulbocastanum* leaf material with about 90% of C16:3(7Z,10Z,13Z)-solamine inhibited the growth of the necrotrophic pathogens *Alternaria solani* and *Botrytis cinerea* (Lenz et al. 2025, accepted).

In the present study, we investigated for the first time potential effects of *S. bulbocastanum* on the planthopper *P. leporinus* in comparison with the commercial potato cultivar ‘Gala’. In no-choice and choice bioassays we investigated both the preferences of the planthopper and its mortality after feeding on the two test plants. We analyzed the content of planthoppers and their gastro-intestines by LC-MS for proof of the cause of their death. Furthermore, the volatile compounds of the wild relative and the potato cultivar were analyzed by GC-MS to find out, if the odors may differ or could act as olfactory cues to the planthopper.

We discuss *S. bulbocastanum* as a potential trap plant for *P. leporinus*, the role of its bioactive compounds and its use in resistance breeding as medium to long-term management options to control *P. leporinus* along with other important potato pests and diseases.

## 2 Material and Methods

### 2.1 No-choice mortality experiments

#### Planthoppers

Two no-choice mortality experiments took place subsequently at the Julius Kühn Institute in Dossenheim, Germany. For the first no-choice experiment, planthoppers were collected on 24 June 2024 in Southwestern Germany with sweeping nets at different sites (potato fields in Bingen 49°56’52.8’’N 7°56’03.6’’E, and Frankenthal 49°33’03.9’’ 8°23’22.1’’E, winter wheat field in Bürstadt 49°38’17.5’’N 8°24’57.6’’E). Likewise, planthoppers were collected in Bad Wimpfen (sugar beet field 49° 13’ 18.7’’N 9° 8’ 36’’E) on 15 July 2024 for the repeated no-choice mortality experiment. The planthoppers have been stored in insect cages and transported to the experiment site. There, ten to eleven individuals have been placed in each of eight transmission cylinders planted with *S. tuberosum* cultivar ‘Gala’, and *S. bulbocastanum*. Specimen of *P. leporinus* only were used after morphological identification (Biedermann, Robert and Niedringhaus, Rolf 2009). The experiments started the same day of the field collection. Both experiments took place in the same climate chamber at 22 °C, 55% relative humidity, and a light/dark cycle of 18/6 h. Insects that died during the experiment were removed daily and stored at −20 °C until further analysis. Same applied to the remaining insects at the end of the trial, i.e. after 11 days.

#### Plant material

*S. tuberosum* cultivar ‘Gala’ (Julius Kühn Institute (JKI), Institute for Breeding Research on Agricultural Crops, Sanitz, Germany) was planted in 3L pots filled with potting soil and sand in a ratio of 40:10 L on 3 June 2024. One tuber per pot was planted. Equal planting applied to *S. bulbocastanum* WKS31741, clonal line blb2G (WKS31741 seed material obtained from Federal ex situ Gene Bank of the Leibniz Institute of Plant Genetics and Crop Plant Research, Gatersleben, Germany), clonal line maintained from JKI (Sanitz, Germany), which resulted in one shoot per pot. All plants were grown in an insect-free greenhouse. The same *S. bulbocastanum* plants were used for the first and the second experiment since this species grew slowly and tuber material was rare. In the case of ‘Gala’, all shoots but one were cut at emerging stage to fit inside the transmission cylinder (Therhaag et al. 2024a) for the first experiment. For the second experiment, a new shoot from the same plant which emerged on 5 July 2025 was taken.

### 2.2 Dual-Choice experiments

Two subsequent experiments were conducted with both plant species in the same climate chamber with equal parameters as described above, starting on 5 and 10 July 2024, respectively, and merged to a choice experiment with a total of 12 replications. One pot with *S. bulbocastanum* and one pot with *S. tuberosum* were put into each of the insect cages (BugDorm 2S120, MegaView Science Co., Ltd., Taiwan). Planthoppers were collected in Southwestern Germany at the same site as described earlier (Bad Wimpfen) on 5 July 2024 and on 9 July 2024. Eight female and two male specimens of *P. leporinus* were placed in the middle of the cages at the same distance to both plants. The number of planthoppers on each plant was counted after 30 minutes, one hour, two hours, three hours, one day and two days later (i.e. after 24 h and 48 h respectively).

### 2.3 Assay for staining plant and insect tissues

Adults of *P. leporinus* from insect rearing on sugar beet variety ‘Danicia’, at the Julius Kühn Institute in Dossenheim, Germany, were taken for the assay of plant sap uptake of *S. bulbocastanum* and *S. tuberosum* cv ‘Gala’.

Cut twigs of potato cultivar ‘Gala’ or *S. bulbocastanum* were inserted through a 1 cm thin foam plate inside Falcon tubes filled with 4 ml solution of 0.5% Brilliant Blue (CAS-No. 3844-45-9, Carl Roth GmbH, Germany). The twig was cut at the bottom and top before insertion to facilitate quick water transport by transpiration. Single individuals of *P. leporinus*, female or male, were transferred to the air-filled part of Falcon tubes. The plate between the solution and the potato leaves prevented the planthopper from falling into the solution.

The tube was covered with a mesh for air circulation and stored at room temperature for two days. After two days, both the plant and the insects were removed from the tube.

For proof of the uptake of blue-dyed water into the xylem of the plants, stems of random samples of each species were cut with razor blades, viewed under a stereo microscope (ZEISS Axioplan) and photographed with a mounted digital camera (Olympus ColorView IIIu FW) using the imaging software analySIS docu 5.0 (Olympus Soft Imaging Solutions GmbH).

The insects were fixed with insect pins on dental wax in a petri dish and dissected in phosphate-buffered solution as described by Takeshita and Kikuchi (2020). The gastro-intestine and its content were easily recognizable by the blue colorant and cut off the insect body ventrally. Each sample was photographed under a stereo microscope (SteREO Discovery.V12, Carl Zeiss Microscopy GmbH, Jena, Germany) equipped with a camera (Axiocam 305, Carl Zeiss Microscopy GmbH, Jena, Germany), then transferred to 10 µl methanol (LC-MS grade) in 1.5 ml reaction tubes and stored at −20 °C until further extraction.

### 2.4 LC/MS metabolite profiling

#### Sample collection and extraction

The LC/MS metabolite profiling of whole insects was obtained from planthoppers of the second no-choice mortality trial. For this, dead planthoppers were collected daily in 1.5 ml reaction tubes and stored at −20 °C. For the LC/MS metabolite profiling of gut extracts, individuals of the assay for plant sap uptake were dissected as described earlier (section 2.3). Crude extracts of whole insects were prepared as follows: A 5-mm steal bead and 50 µl 80% (v/v) methanol/water with 0.5% (v/v) formic acid were added to each insect. The sample was vigorously shaken for 5 min at 30 Hz and room temperature using a mixer mill (MM300, Retsch GmbH, Haan, Germany). After short centrifugation for sample precipitation, samples were treated with ultrasonic at room temperature for 5 min and then vigorously shaken for 5 min at 30 Hz again. After 15 minutes of centrifugation (13000 g, 22 °C), the supernatant was collected for LC-MS analysis. Crude extracts of insect intestinal samples were extracted by adding 20 µl 80% methanol/water with 0.5% formic acid. Samples tissues were not disrupted, only the intestinal content was targeted for analysis. For this, samples were shaken on a vertical rotary shaker for 10 min (1000 rpm, RT) and centrifuged for 10 min (13000 rpm, 22 °C). The supernatant was subjected to LC-MS analysis.

#### LC/MS

Extracts were analyzed on an Infinity 1290 series UHPLC system hyphenated to a 6550 iFunnel Q-TOF mass spectrometer (Agilent Technologies, Santa Clara, CA) via a dual jet stream electrospray ion source at the Julius Kühn Institute in Berlin, Germany as described (Tais et al. 2021). Samples (injection volume 5 µL for whole insect analyses; 10 µl for insect intestinal analyses) were separated on an ACQUITY UPLC HSS T3 column (2.1 mm × 100 mm, particle size 1.8 µm, pore size 100 Å, Waters Corporation, Milford, MA) using 0.1% (v/v) formic acid in water and 0.1% formic acid in acetonitrile as eluent A and B, respectively. The binary gradient program at a flow rate of 500 µL/min was 0-1 min: isocratic 5% B, 1-15 min: linear to 60% B, 15-18 min: linear to 95% B, 18-19 min: isocratic 95% B, 19-21 min: isocratic 5% B. The column temperature was 40 °C; the autosampler temperature 6 °C. Eluting compounds were detected in positive mode from m/z 50-1700 (centroid mode, 3 spectra/s) in all samples and in negative ion mode in intact insect total extracts; instrument settings were applied as described by Tais et al. (2021).

#### Analysis of LC/MS data

Untargeted metabolite profiling was performed using the R package “xcms” Release (3.20) with peak picking using the ‘centWave’ method with a signal to noise ratio of SN = 4, otherwise using default settings. Peaks were aligned without the need of retention time correction. ‘fillPeaks’ was neglected. All peaks, including potential adducts and isotopes from retention times between 0.6 to 14 min were included in statistics. Data were imported in MetaboAnalyst (Xia et al. 2009; Pang et al. 2021, 2022) for visualization and statistical analysis. Raw data inspection, compound spectrum extraction, compound database search and generation of molecular formulas were done using MassHunter Qualitative Analysis software (version B.07.00, Agilent Technologies); in-house libraries (Gorzolka et al. 2025) and the technology platform “Metlin” (Guijas et al. 2018) were used for compound annotation by exact m/z match (5 ppm) as well as retention time match for positive library hits.

### 2.5 GC/MS analysis of Volatile Organic Compounds (VOCs)

#### Collection of VOCs of the potato cultivar ‘Gala’ and *S. bulbocastanum*

VOCs of six plants of the potato cultivar ‘Gala’ and of six plants of *S. bulbocastanum* were collected on 30 July 2024 from plants of the second no-choice mortality experiment. For the collection, a single shoot of each sample plant was placed inside a plastic oven bag which was heated beforehand at 60 °C for 4 hours and cooled again. Each bag was connected to a headspace collection device (FLUSYS GmbH, Offenbach, Germany), which was developed by J. Gross (Rid et al. 2016; Gross et al. 2019) and improved in the PurPest Project (European Commission 2022) described by Karimi and Gross (2024). The collection was implemented in an insect-proof greenhouse chamber. The VOCs were trapped in stainless steel tubes filled with Tenax TA35/60 sorbent (Markes, Neu-Isenburg, Germany) until further analysis. The air inside of empty plastic bags was collected and analyzed as a control to avoid identification of compounds not originating from the plants.

#### Analysis of GC/MS data

A subsequent chemical analysis of the VOCs was undertaken at the day of collection through automated thermodesorption (TurboMatrix™ ATD 650, PerkinElmer, Rodgau, Germany) and gas chromatography coupled with mass spectrometry at the Julius Kühn Institute in Dossenheim, Germany as described by Gallinger et al. (2020). In our study, the VOC flow was split so that 30% of the flow were subjected to a full-scan mass spectrometer monitoring at a range of 50-350 m/z. Obtained data were analyzed with AMDIS software (Automated Mass spectral Deconvolution and Identification System, V. 2.71; National Institute of Standards and Technology NIST, Boulder, CO) according to a protocol by Gross et al. (2019). In AMDIS, ion fragmentation patterns, retention times and retention indices (RI) were compared for the identification of the detected compounds with those of standard compounds that were introduced in the same system and stored in an in-house library. For compounds not included in the in-house library, matches of the NIST library (National Institute of Standards and Technology) and the NIST Chemistry WebBook (2025) were taken for further comparison. In that case, only RI from gas chromatography systems with similar technical parameters in terms of column diameter and length, stationary phase material and comparable temperatures were considered. Compounds identified with the use of authentic standards are treated as “identified”, those obtained without the use of authentic standards as “annotated”. Compounds that could not be identified are annotated as “unknown”. The identification criteria of the AMDIS software were as follows: the minimum match factor was set to 80% using retention index (RI) data, the RI window was set to 20 with the match factor penalties set at very low level with 20 for maximum penalty and 20 for “no RI in library”. Compounds occurring in less than two (out of twelve) samples were excluded from the analysis as well as the internal standard and known synthetic compounds. Relative proportions of each VOC were calculated by division of the peak area through the sum of the peak areas of all compounds per sample.

### 2.6 Statistical analysis

Statistical analysis of the mortality experiments, the choice-experiments and the analysis of VOC profiles was performed in RStudio/2024.12.0+467 (“Kousa Dogwood” Release for windows, Posit Software, PBC) using R version 4.4.2 (R Core Team 2024. R: A Language and Environment for Statistical Computing. R Foundation for Statistical Computing, Vienna, Austria. https://www.R-project.org/). In case of the no-choice mortality experiments, a multivariate Cox regression analysis was performed using the coxph() function of the R ‘survival’ package (Therneau et al. 2024) considering ‘treatment’, i.e. the two different plant species, and ‘pot’ as covariates to account for the grouping of individuals in the pots with the different plant species. Survival curves were created with the survfit() function of the same package.

With regards to the choice experiments investigating plant preferences, the counted number of planthoppers per plant species in each cage was analyzed using a generalized linear mixed model approach with a zero-inflated Poisson distribution and log-link function of the ‘glmmTMB’ package (Brooks et al. 2017; McGillycuddy et al. 2025). The number of individuals still alive at the time of counting was taken into account as an offset. The model was fitted with plant species as fixed effect, the cages and observation timings as random effects. Model fit parameters were obtained from the summary() function. An evaluation of the differences, i.e. the contrasts, between the marginal means of the contrasting variable ‘plant species’ at the different observation timings was conducted using the averaged contrast analysis of the ‘modelbased’ package (Makowski et al. 2025) with the function estimate_contrasts(). The confidence interval level was set to 0.95, and the Holm-Bonferroni method for p-adjustments in frequentist multiple comparisons was applied. Results were visualized using the ‘ggplot2’ package in R (Wickham 2016).

For the VOC profiles of the both species, a non-metric multidimensional scaling using Bray-Curtis dissimilarities plot was obtained using metaNMDS from the ‘vegan’ package in R (Brückner and Heethoff 2016; Oksanen et al. 2025). Of the same package, the envfit() function was applied to the compositional data set with permutations set to 10,000. Anderson’s permutational multivariate analysis of variance and of dispersions (Anderson 2001; Anderson et al. 2006) has been computed in R with PERMANOVA and PERMDISP from the Bray-Curtis dissimilarity matrix of the VOCs as described in Brückner and Heethoff (2016).

## 3 Results

### 3.1 No-choice mortality experiments

Two no-choice experiments were conducted with field-caught adults of *P. leporinus* which were placed inside cylinders with either *S. tuberosum* cv ‘Gala’ or *S. bulbocastanum* plants. For a period of 11 days, dead planthoppers were counted and collected daily. Analysis of mortality was undertaken with the number of dead planthoppers per day and plant species.

Mortality increased significantly for planthoppers feeding on *S. bulbocastanum* revealed by several distinct significance tests (Table 1). Both experiments showed that the effect of the grouping in ‘pot’ fails significance whereas the effect of the ‘treatment’, namely the effect of the different plant species exposed to the planthoppers, contributes significantly to the hazard ratio of death of the planthoppers of the studies. Survival plots of the planthoppers of both no-choice experiments are compared in Figure 1 and visualize impact of the plant species on the survival.

**Table 1.** Results of statistical tests of a multivariate Cox regression analysis of both no-choice experiments. The covariate ‘treatment’ refers to the two different plant species, i.e. *S. bulbocastanum* and the potato cv ‘Gala’, whereas the covariate ‘pot’ accounts for the grouping of the planthoppers to the plant species potted in 16 and 12 pots in experiment I (collected from potato and winter wheat) and II (collected from sugar beet), respectively. Significance codes apply as follows: ‘***’ for p < 0.001, ‘**’ for p < 0.01, ‘*’ for p < 0.05. se: standard error, z: z-value, df: degrees of freedom. The Pr(>|z|) column represents the p-value associated with the value in the z value column.

|  | *Analysis of experiment I* |  | *Analysis of experiment II* |  |
| --- | --- | --- | --- | --- |
| **Concordance** | 0.779 | se = 0.018 | 0.665 | se = 0.039 |
| **Likelihood ratio test** | 121.2 on 2 df | $p = < 2e-16$ | 22.77 on 2 df | $p = 1e-05$ |
| **Wald test** | 96.75 on 2 df | $p = < 2e-16$ | 22.5 on 2 df | $p = 1e-05$ |
| **Score (logrank) test** | 128.6 on 2 df | $p = < 2e-16$ | 23.93 on 2 df | $p = 6e-06$ |

| *Analysis of experiment I* |  |  |  |  |  |
| --- | --- | --- | --- | --- | --- |
| *Covariate* | *coef* | *exp(coef)* | *se(coef)* | *z* | *Pr(> \|z\| )* |
| ‘treatment’ | -2.93991 | 0.05287 | 0.420099 | -6.983 | 2.88e-12 *** |
| ‘pot’ | 0.05088 | 1.05220 | 0.04090 | 1.244 | 0.214 |

| *Analysis of experiment II* |  |  |  |  |  |
| --- | --- | --- | --- | --- | --- |
| ‘treatment’ | -1.04751 | 0.35081 | 0.43181 | -2.426 | 0.0153 * |
| ‘pot’ | 0.00998 | 1.01003 | 0.06180 | 0.162 | 0.8717 |

**Fig. 1.**
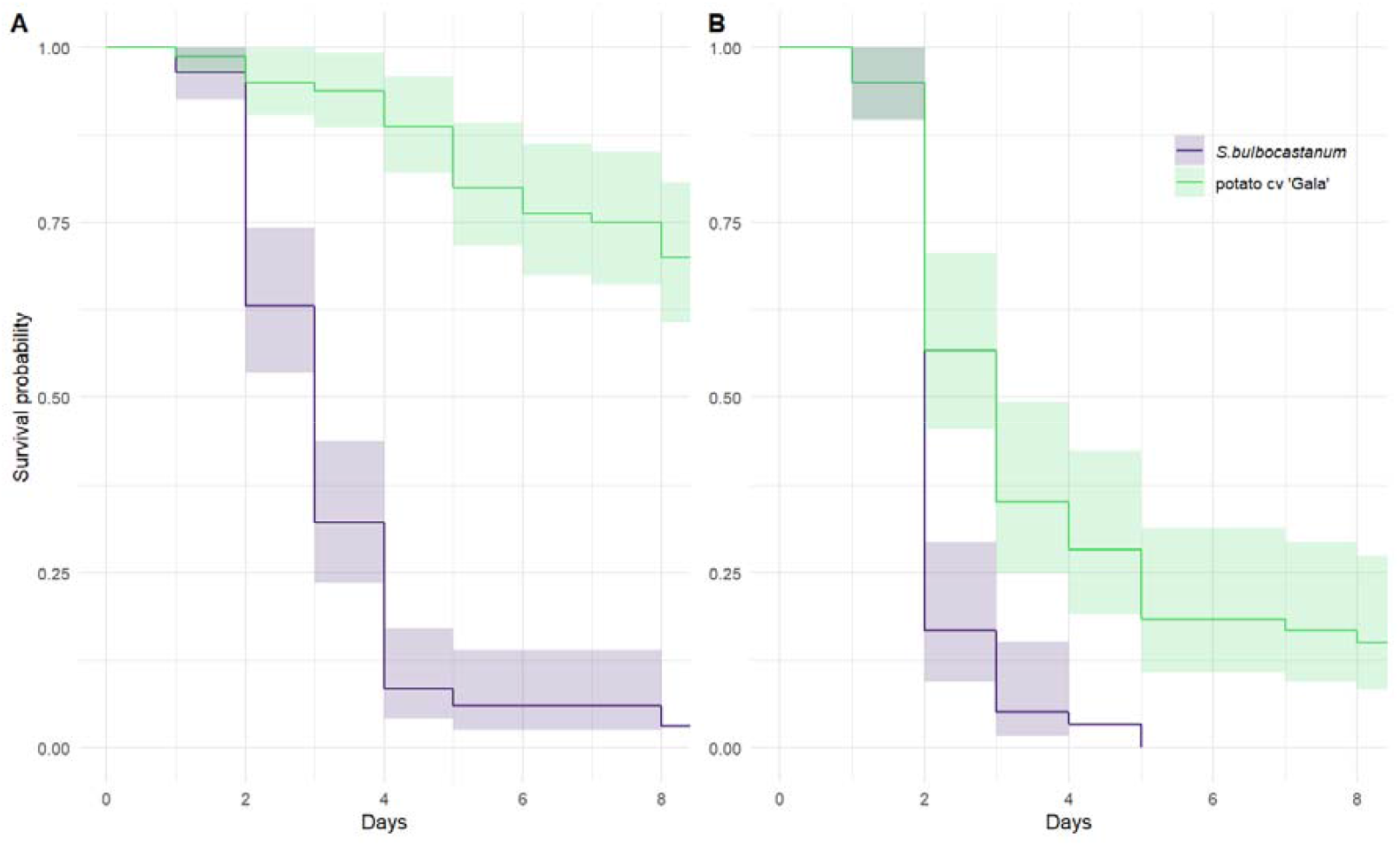
Survival plots including the confidence interval (0.95) of (A) the first (adults collected from potato and winter wheat), and (B) the second (adults collected from sugar beet) no-choice experiment (B) with *P. leporinus*

### 3.2 Dual-Choice experiment

In the dual-choice experiment, field-caught planthoppers were placed inside the insect cages and their total number was counted on each plant species at different observation times over a period of two days, with individuals still alive chosen as offset.

As can be drawn from Figure 2, planthoppers which were decisive for a plant, preferred the crop wild relative at the different observation timings. A marginal contrast analysis with a Holm-Bonferroni method for p-adjustments revealed significance levels ranging between p < 0.001 and p < 0.05 (Table 2) in the chosen confidence interval of 95%.

**Table 2.**
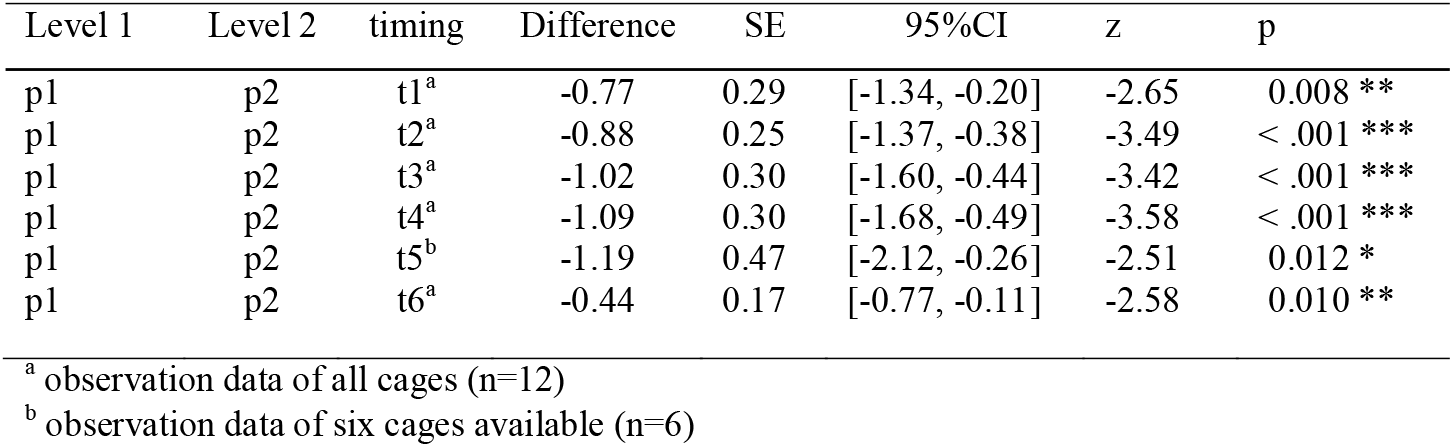
Results of the average contrast analysis of the choice experiment displaying the differences of the marginal means of the contrasting variable ‘plant species’ at different observation timings, with p1: *S. bulbocastanum*, p2: *S. tuberosum* cv ‘Gala, SE: standard error, z: z-value, df: degrees of freedom, and t1 to t6: 0.5h, 1h, 2h, 3h, 24h, 48h. P-values are obtained from the Holm-Bonferroni method with significance codes as follows: ‘*’ for p < 0.05, ‘**’ for p < 0.01, and ‘***’ for p < 0.001 within the 95% confidence interval (95% CI)

**Fig. 2.**
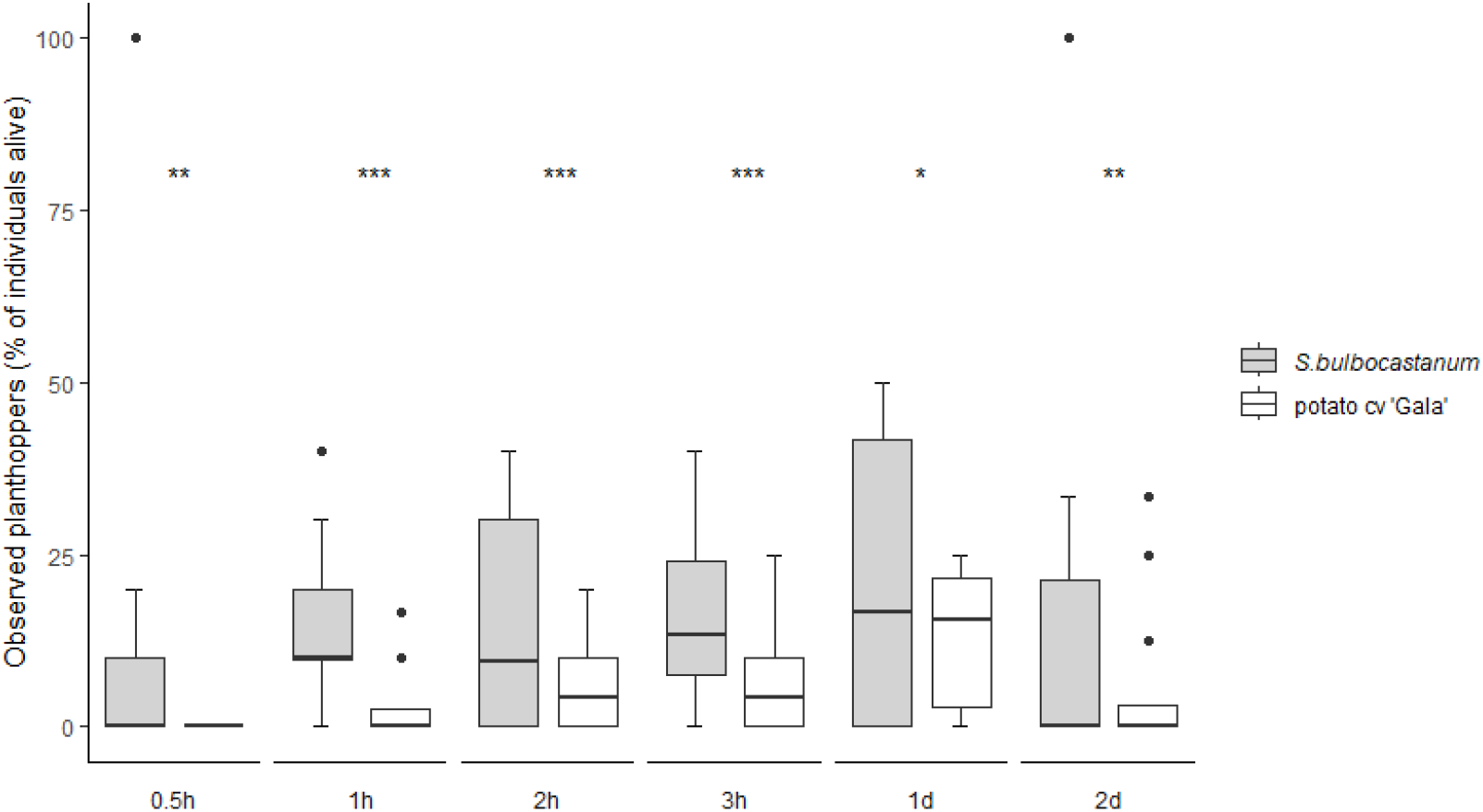
Preference of planthoppers for a plant species *S. bulbocastanum* or *S. tuberosum* cv ‘Gala’ observed in cages (*n*=12) of the choice experiment. Boxplots represent the percentage of observed planthoppers related to the number of planthoppers still alive; bars indicate standard error (SE); at the observation timing 1d, only data of six cages (*n*=6) are available; asterisks indicate significant adjusted p-values obtained from the Holm-Bonferroni method with ‘*’ for p < 0.05, ‘**’ for p < 0.01, and ‘***’ for p < 0.001, in the 95% confidence interval

### 3.3 Assay for plant sap uptake

For the LC/MS analysis of planthopper intestines, blue dyed water was used for plant and eventually insect uptake to facilitate recognition and excision of the intestines. Photographs of petiole sections show some cells with blue content, presumably xylem cells stained by uptake of blue colored water by transpiration (Figure 3 A). Photographs of dissected insects show the blue stained intestine could be easily distinguished from other organs (Figure 3 B).

**Fig. 3.**
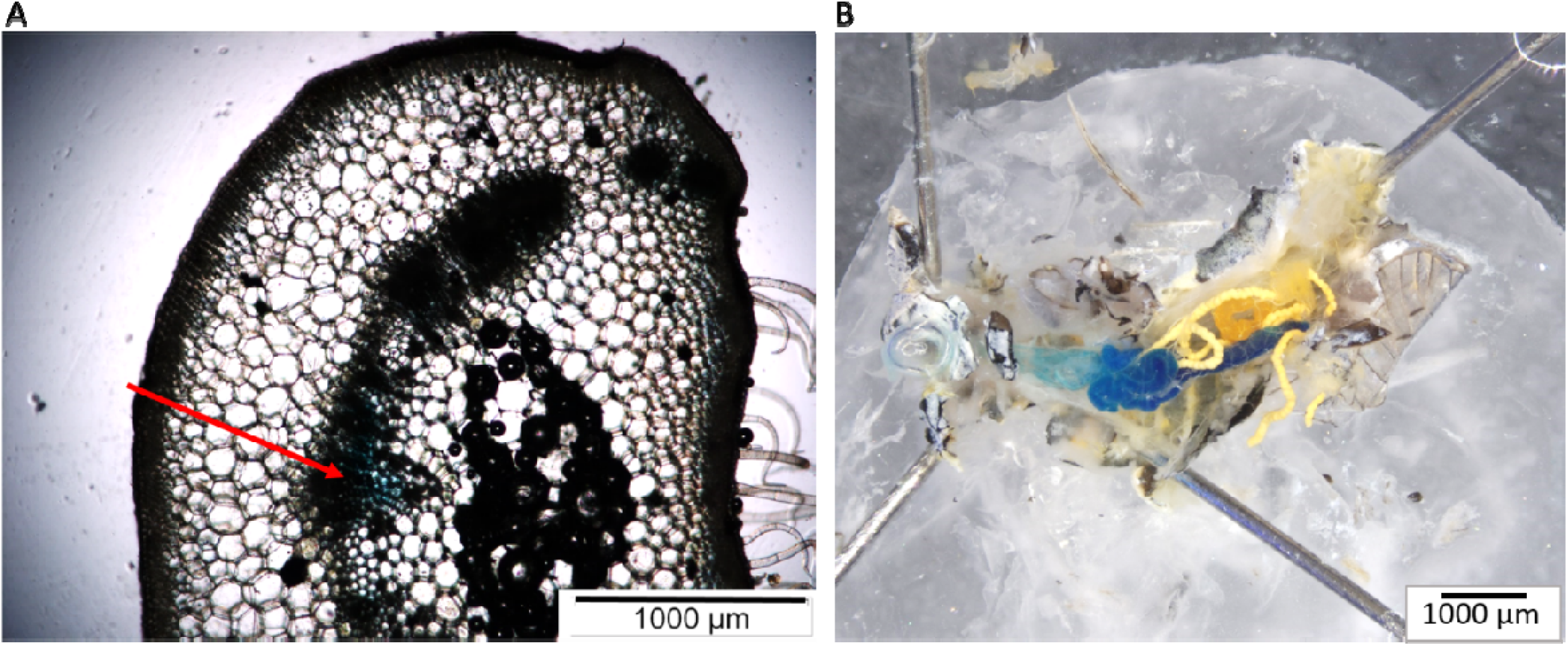
A: Photograph of a section of a cut potato leaf petiole after insertion of the twig in blue-dyed water for two days (25x magnification); red arrow points to cells of the vascular bundle filled blue. B: Stacked photograph before excision of intestine with blue content of female *P. leporinus*, 15.2x magnified

### 3.4 LC/MS analysis

The untargeted LC/MS-analysis of a) the extracts of the whole insect bodies, and b) the extracts of intestines of the planthoppers resulted in 2,544 and 1,976 signals with retention times between 0.6 min to 14 min, respectively. A fivefold-change analysis was calculated to compare the absolute value of change between the two group means of planthoppers which fed on *S. bulbocastanum* vs. *S. tuberosum* cv ‘Gala’. An adjusted p-value (FDR) threshold of p<0.05 was taken for the unpaired t-tests. Figure 4 shows a volcano plot combining the fold change analysis and the t-test values of the detected compounds with the acyl solamine signals labelled. This analysis reveals that out of the 2,544 analyzed signals in the insect bodies, only C16:3-solamine [M+H]+ was significantly more than 5-fold enhanced in insects that fed on *S. bulbocastanum* with p<0.05. Of the 1,976 detected compounds of the intestine extracts, also acyl solamine signals (C16:3-solamine [M+H]+, C16:3-solamine [M+H]2+) had the highest fold change enhancement and significance level in *S. bulbocastanum* derived planthoppers. The significantly changing metabolites of both datasets are listed in Table 3.

**Table 3.** Fold change and significance analyses of metabolite profiling data obtained from LC-MS extracts of A) whole plant hopper with in total 2,544 signals and B) plant hopper intestines with in total 1,976 signals. Thresholds were 5-fold change (FC) and FDR-adjusted p-values of p<0.05. Identified compounds are indicated in “annotation”. Signals “Unknown 3 Isotope 1-3” derived from one single metabolite. C16:3-sol: C16:3-solamine. A fold change of < 1 indicates enhanced abundances in *Solanum bulbocastanum* derived plant hopper samples, an FC value of > 1 indicates enhanced abundances in *Solanum tuberosum* derived plant hopper samples. “Unknown”: no match in data bases

|  | signal | annotation | FC | log2(FC) | p.adjusted | -log10(p) |
| --- | --- | --- | --- | --- | --- | --- |
| A | 8.77_448.4253 | C16:3-sol [M+H]<sup>+</sup> | 0.040354 | -4.6311 | 1.19E-17 | 16.924 |
|  | 12.19_268.2742 | Unknown 1 | 9.3015 | 3.2175 | 1.34E-09 | 8.8716 |
|  | 12.90_491.3095 | Unknown 2 | 8.6235 | 3.1083 | 2.27E-06 | 5.6447 |
| B | 8.80_448.419 | C16:3-sol [M+H]<sup>+</sup> | 0.0012672 | -9.6241 | 4.30E-11 | 10.366 |
|  | 8.80_224.713 | C16:3-sol [M+2H]<sup>2+</sup> | 0.0014057 | -9.4745 | 4.63E-11 | 10.335 |
|  | 8.34_581.151 | Unknown 3 Isotope 3 | 0.030953 | -5.0138 | 0.0056315 | 2.2494 |
|  | 8.34_579.150 | Unknown 3 Isotope 1 | 0.11415 | -3.1311 | 0.0068515 | 2.1642 |
|  | 8.34_580.154 | Unknown 3 Isotope 2 | 0.11547 | -3.1144 | 0.0077163 | 2.1126 |
|  | 13.53_454.283 | Unknown 4 | 0.1149 | -3.1216 | 0.036768 | 1.4345 |

**Fig. 4.**
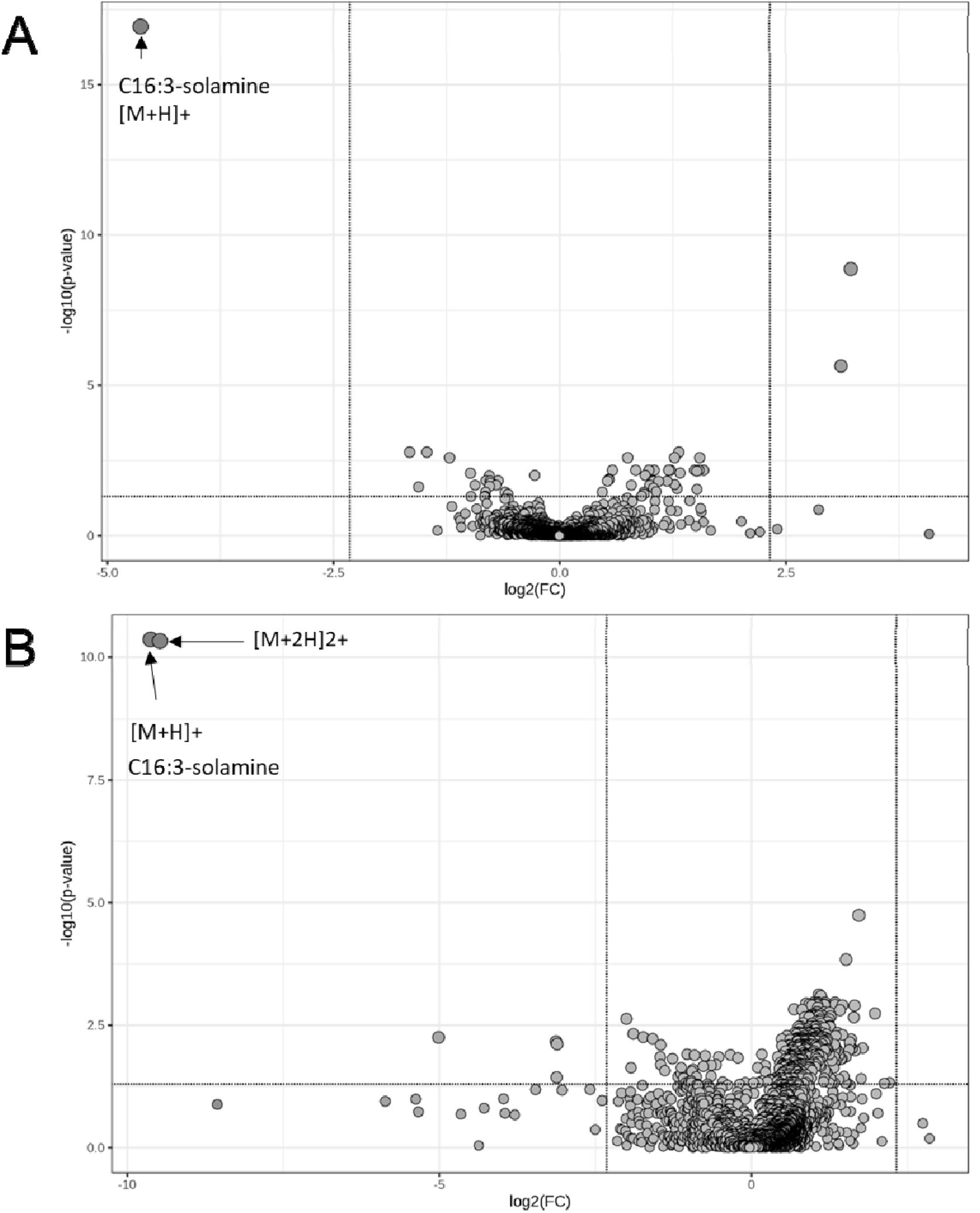
Volcano plots of untargeted metabolite profiling data with unpaired samples of extracts of whole insect bodies of *P. leporinus* (A) and extracts of intestines from planthoppers (B) fed on *S. bulbocastanum* vs. *S. tuberosum* cv ‘Gala’. The x-axis shows the fold change (FC) of the compound in log2 scale, the y-axis shows the significance of the fold change as −log10 (p-value) based on FDR adjusted p-values from the t-tests. Points indicate compounds; compounds enhanced in *S bulbocastanum* group on the left side, compounds enhanced in *S. tuberosum* cv ‘Gala’ on the right side. Vertical solid lines indicate thresholds of 5x fold change. Horizontal lines indicate a significance threshold of p=0.05, points above this line are significantly (p<0.05) enhanced in one of the test groups. Arrows indicate C16:3-solamine as single charged ion [M+H]+ and detected double charged ion [M+2H]2+. Table 3 presents all compounds with >5 fold change with p<0.05

A Mann-Whitney U test was applied to the data of the C16:3-solamine peak area of the LC/MS analyses of extracts obtained from planthoppers either feeding on *S. bulbocastanum* or *S. tuberosum* cv ‘Gala’. With regards to extracts of whole planthoppers, a significance level of p < 0.00001 was revealed and C16:3-solamine was undetectable in individuals feeding on *Solanum tuberosum* ‘Gala’ (Figure 5). In intestines of planthoppers, a significant difference of this compound was observed with a p-value of 0.001. Differentiation by the sex of the planthoppers demonstrated a significant uptake of the C16:3 solamine at p < 0.001 of the male and female planthoppers fed on *S. bulbocastanum*, thus no sex-specific differences in the feeding behavior of plant hoppers were observed (Figure 6).

**Fig. 5.**
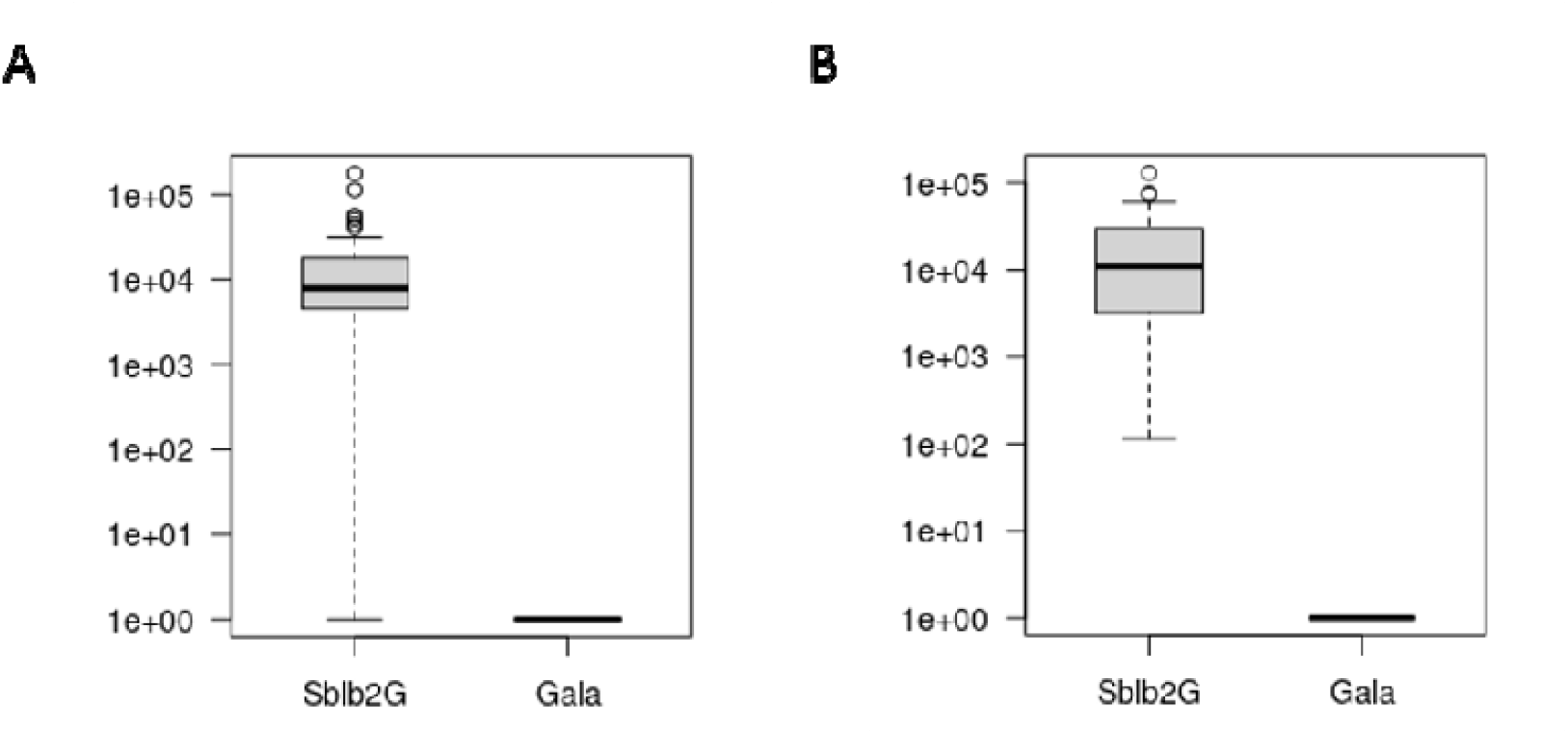
Boxplots of the C16:3-solamine peak area in extracts of whole planthopper bodies (A) and extracts of intestines (B) of planthoppers after feeding on *Solanum bulbocastanum* vs. *Solanum tuberosum* cv ‘Gala’. The x-axis indicates the two plant species (here, Sblb2G: S. *bulbocastanum*, Gala: *S. tuberosum* cv ‘Gala’), the y-axis indicates the Log10 peak area of the LC/MS data

**Fig. 6.**
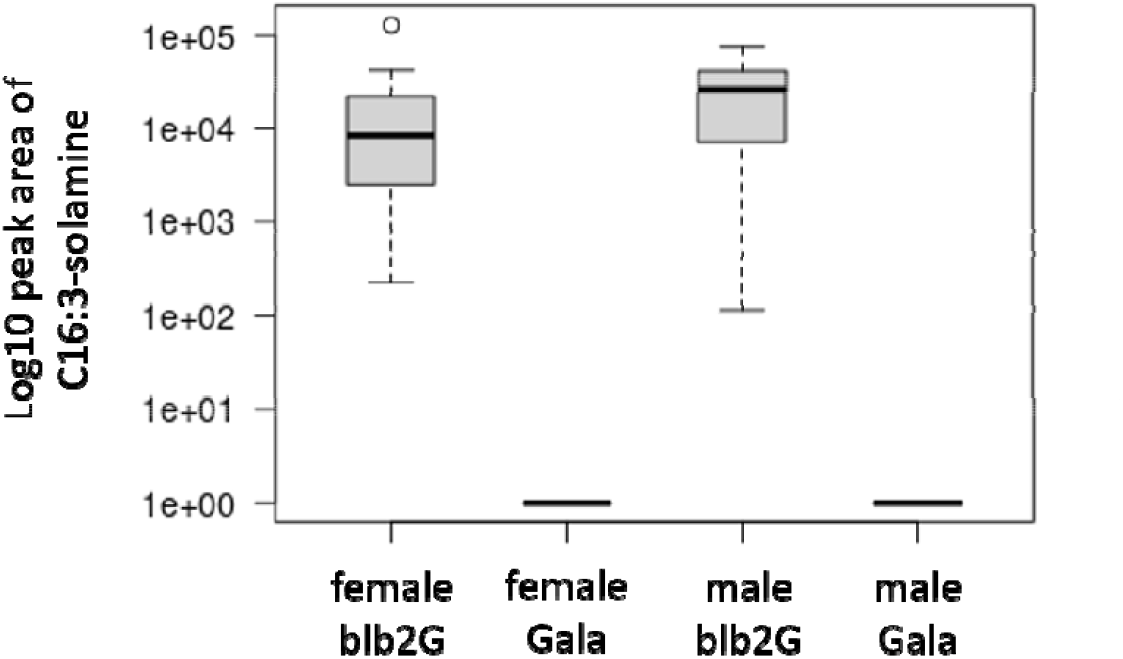
Boxplots of the C16:3-solamine peak area in extracts of intestines of female and male planthoppers after feeding on *Solanum bulbocastanum* vs. *Solanum tuberosum* cv ‘Gala’. The x-axis indicates the two sexes (female *P. leporinus*, male *P. leporinus*) fed on the two plant species (here, blb2G: *S. bulbocastanum*, Gala: *S. tuberosum* cv ‘Gala’), the y-axis indicates the Log10 peak area of the LCMS data of the C16:3-solamine

### 3.5 GC/MS analysis

In total, 80 compounds of the collected VOCs of *S. bulbocastanum* and of *S. tuberosum* cv ‘Gala’ resulted after selection as described in the section ‘Material and Methods’. A table with all compounds is provided in Online Resource 1 (ESM_1.pdf). Thereof, 32 compounds were found in ‘Gala’ only and seven compounds only in the wild potato. These differentiating compounds are listed in Table 4. Chromatograms of the GC/MS data of the collected VOCs show the very distinct profiles of the two species (Figure 7). Compounds which contribute significantly (p<0.001) to a differentiation of the VOC patterns are inserted as arrows in a non-metric multidimensional scaling plot (Figure 8).

**Table 4.** List of identified, annotated or unknown volatile organic compounds only found in *S. bulbocastanum* or *S. tuberosum* cv ‘Gala’, indicating the retention indices (RI), p-values and species. Compounds differentiating the VOC pattern after application of environmental vector analysis with a permutation of 10,000 are listed with following significance codes: ‘***’ for p < 0.001, ‘**’ for p < 0.01, ‘*’ for p < 0.05

| Compound | RI | Pr(>r) | species |
| --- | --- | --- | --- |
| 3-Hexen-1-ol, (E)-<sup>a</sup> | 852 | 0.264174 | *S. bulbocastanum* |
| 2-Phenylethanol<sup>b</sup> | 1112 | 0.750825 | “ |
| unknown_RI_1363 ($m/z$ 43, 57, 41, 70, 71) | 1363 | 0.711729 | “ |
| 1-Tetradecene<sup>a</sup> | 1392 | 0.001600 ** | “ |
| unknown_RI_1458 ($m/z$ 165, 193, 151, 95, 208) | 1458 | 0.207779 | “ |
| unknown_RI_1496 ($m/z$ 71, 57, 43, 85, 41) | 1496 | 0.694931 | “ |
| unknown_RI_1543 ($m/z$ 55, 54, 71, 43, 41) | 1543 | 0.496550 | “ |
| α-Pinene<sup>b</sup> | 932 | 0.516648 | *S. tuberosum* cv ‘Gala’ |
| β-Myrcene<sup>b</sup> | 991 | 0.010999 * | “ |
| α-Terpinene<sup>a</sup> | 1015 | 0.188081 | “ |
| (R)-(+)-Limonene <sup>b</sup> | 1027 | 0.008999 ** | “ |
| unknown_RI_1043 (*m/z* 87, 102, 57, 41, 130) | 1043 | 0.006799 ** | “ |
| 4,8-dimethyl-1(E)3,7-nonatriene <sup>b</sup> | 1117 | 0.066993 | “ |
| Benzothiazol <sup>b</sup> | 1222 | 0.023298 * | “ |
| unknown_RI_1339 (*m/z* 121, 93, 107, 79, 41) | 1339 | 0.001700 ** | “ |
| α-Cubebene <sup>a</sup> | 1351 | 0.003900 ** | “ |
| (-)-α-Copaene <sup>a</sup> | 1378 | 0.042496 * | “ |
| (-)-β-Bourbonene <sup>a</sup> | 1387 | 0.003800 ** | “ |
| α-Gurjunene <sup>a</sup> | 1412 | 0.004500 ** | “ |
| β-Caryophyllene <sup>b</sup> | 1424 | 0.001700 ** | “ |
| trans-α-Bergamotene <sup>a</sup> | 1438 | 0.033397 * | “ |
| unknown_RI_1445 (*m/z* 69, 41, 91, 55, 77) | 1445 | 0.011199 * | “ |
| (E)-β-Farnesene <sup>a</sup> | 1459 | 0.003500 ** | “ |
| cis-Muurolo-4(15),5-diene <sup>a</sup> | 1466 | 0.002700 ** | “ |
| γ-Muuroloene <sup>a</sup> | 1480 | 0.002200 ** | “ |
| (-)-Germacrene D <sup>a</sup> | 1485 | 0.001300 ** | “ |
| α-Farnesene <sup>a</sup> | 1509 | 0.262674 | “ |
| γ-Cadinene <sup>a</sup> | 1518 | 0.002800 ** | “ |
| δ-Cadinene <sup>a</sup> | 1527 | 0.170583 | “ |
| α-Calacorene <sup>a</sup> | 1547 | 0.004000 ** | “ |
| unknown_RI_1572 (*m/z* 105, 67, 111, 122, 41) | 1572 | 0.002300 ** | “ |
| Caryophyllene oxide <sup>a</sup> | 1587 | 0.216278 | “ |
| unknown_RI_1608 (*m/z* 41, 67, 82, 111, 71) | 1608 | 0.011899 * | “ |
| α-Corocalene <sup>a</sup> | 1626 | 0.017798 * | “ |
| τ-Cadinol <sup>a</sup> | 1645 | 0.002000 ** | “ |
| unknown_RI_1679 (*m/z* 183, 198, 168, 153, 165) | 1679 | 0.003700 ** | “ |
| unknown_RI_1734 (*m/z* 43, 91, 79, 121, 55) | 1734 | 0.002600 ** | “ |
| unknown_RI_1844 (*m/z* 69, 41, 81, 55, 93) | 1844 | 0.168583 | “ |
| unknown_RI_1972 (*m/z* 69, 93, 41, 81, 107) | 1972 | 0.155984 | “ |

<sup>a</sup> Annotated compounds derived from NIST Mass Spectral Search Program and comparison of retention index (RI ) data from NIST Chemical WebBook
<sup>b</sup> Compounds identified by comparing mass spectrum and RI with authentic standards on the same GC-MS Mass fragments for unknowns are listed with the base peak and other fragments in order of decreasing abundance

**Fig. 7.**
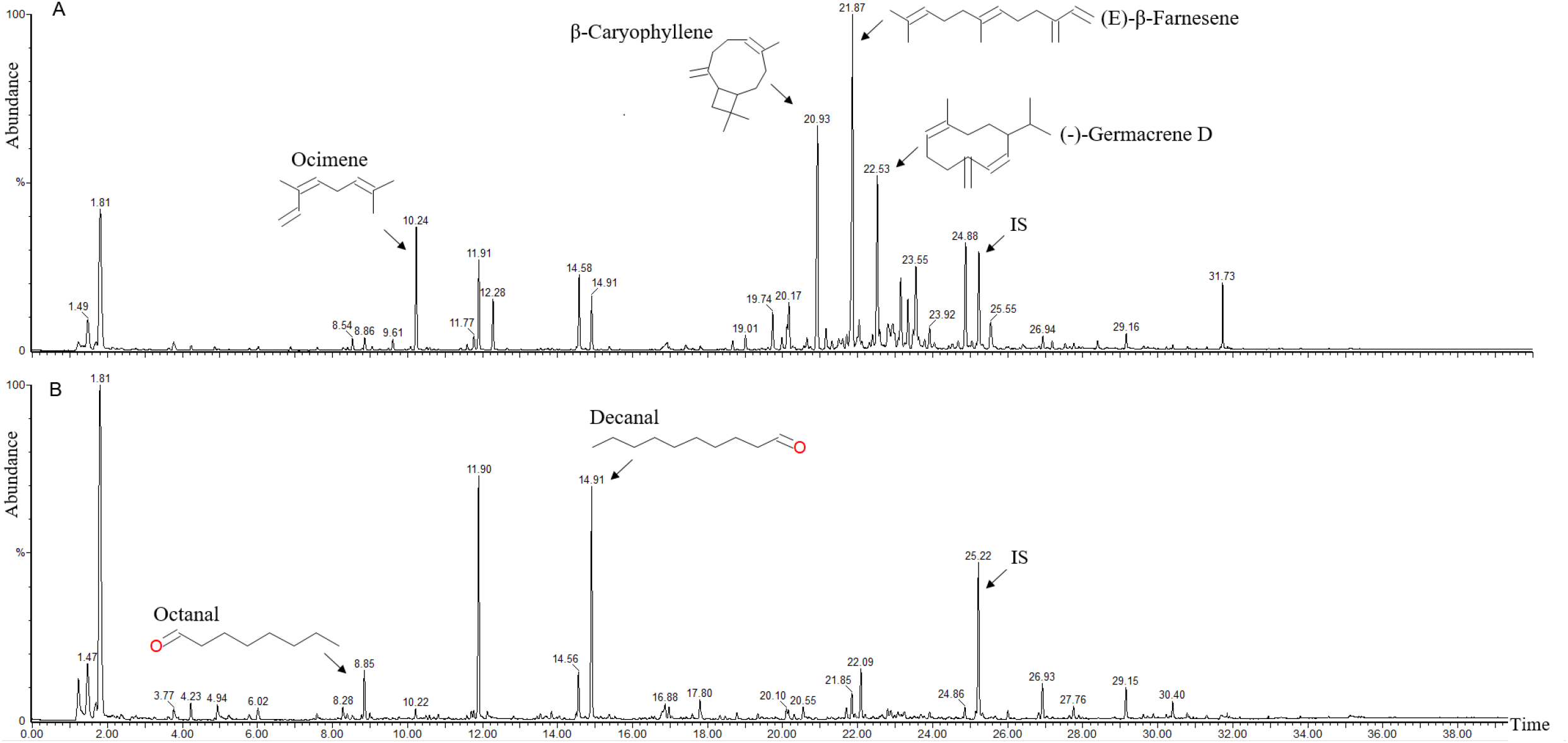
Total ion current chromatograms of volatile organic compounds of *S. tuberosum* cv ‘Gala’ (A) and *S. bulbocastanum* (B) with arrows pointing to selected compounds; dodecanoic acid, ethyl ester was used as internal standard (IS)

**Fig. 8.**
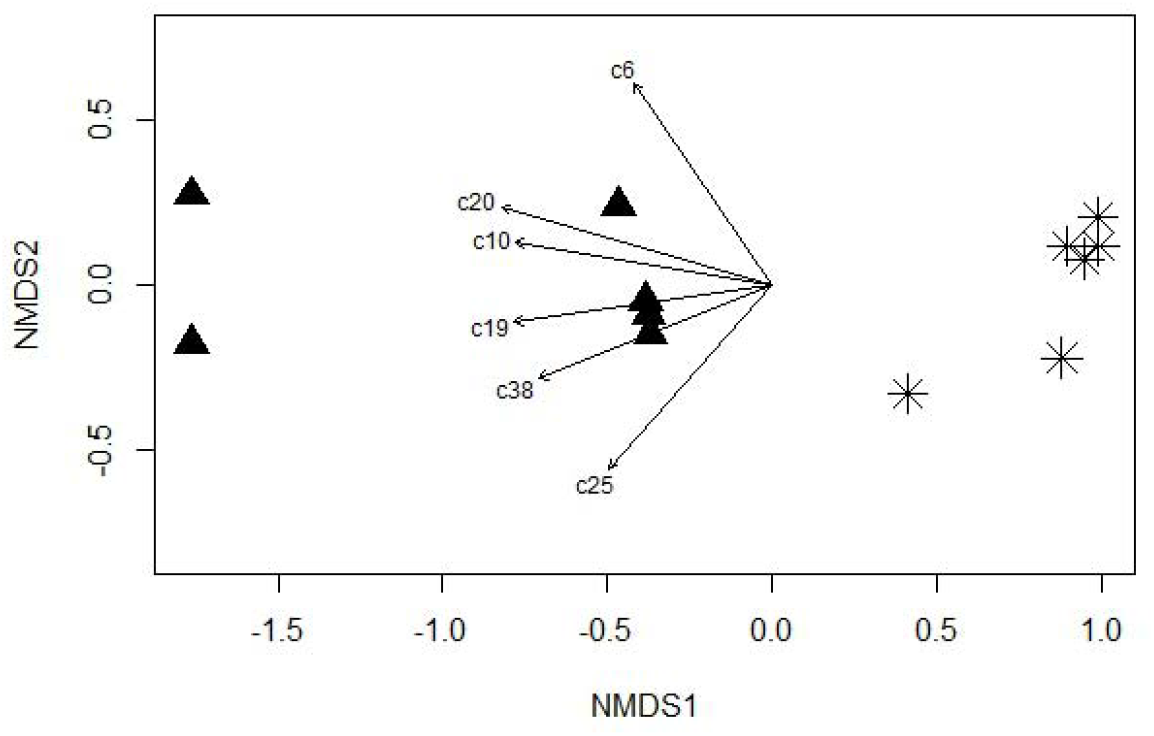
Non-metric multidimensional scaling (NMDS) plot classifying the VOC profiles of *S. bulbocastanum* and *S. tuberosum* cv ‘Gala’ based on Bray-Curtis dissimilarities, displayed as triangles and asterisks, respectively. Compounds (c_i_) accounting for a differentiation of the VOC pattern (with p-values < 0.001) after vector analysis are displayed by arrows. A numbered list of compounds is provided in Online Resource 1 (ESM_1.pdf). The significance of fitted vectors is assessed using permutation (level 10,000) of the 80 target compounds, the goodness of fit statistic is squared correlation coefficient (r^2^)

As shown by the results of the multivariate statistical test PERMANOVA (Table 5), the difference of the two species clearly affects the VOC profiles (with a p-value of 0.0019). It accounts for 70.4 % (R^2^ = 0.704) of the variation in the composition. The PERMDISP test results in a p-value of 0.7133, thus showing no evidence of significant difference in dispersion between the two species. This agrees with the visualization in Figure 8.

**Table 5.** Results of multivariate statistical tests PERMANOVA and PERMSDISP based on Bray-Curtis dissimilarities for the compositional data set. The number of permutations was 10,000

| PERMANOVA |  | PERMDISP |  |
| --- | --- | --- | --- |
| F | Pr(>F) | F | Pr(>F) |
| 23.76 | 0.0019 ** | 0.0761 | 0.7133 |

## Discussion

For the first time plant-insect-interactions between the vector *P. leporinus* and the wild potato *S. bulbocastanum* have been investigated. Mortality experiments designed as no-choice experiments revealed lower survival rates of the vector compared to survival rates on a common potato variety. Since these first results seemed astonishing, subsequent experiments followed, including dual-choice-experiments to investigate whether *S. bulbocastanum* either elicits attraction or rather evasive behavior of the planthopper. The choice-experiments revealed a preference of the planthoppers towards the wild potato rather than the potato cultivar. A study of Gorzolka et al. (2025) discovered a long chain fatty acyl (LCFA) solamine (C16:3(7*Z*,10*Z*,13*Z*)-solamine), which was isolated from the leaves of *S. bulbocastanum*. Its role in plant defense was investigated with interesting results on common potato pests such as bioactivity against the potato flea beetle and the Colorado potato beetle (CPB). Unlike *P. leporinus*, the CPB is not a plant sap sucking insect but feeds on leaf tissue.

Likewise, the potato flea beetle feeds on leaf tissue pitting little holes into potato leaves. In feeding no-choice assays, CPB fed significantly less leaf material of *Solanum bulbocastanum* compared to the *Solanum tuberosum* varieties ‘Maxi’ and ‘Quarta’. Small quantities of exogenous applied C16:3-solamine led to significant lower feeding activity with > 70% less leaf consumption for the CPB and > 80% for the potato flea beetle. However, *S. bulbocastanum* leaf tissue or the exogenous applied C16:3-solamine were not sufficient to lead to acute toxic effects on these pests (Gorzolka et al. 2025). Compared to the no-choice test of our study, we did not study the feeding activity of the planthoppers but their survival rate. However, the planthoppers sucked on both, the stalks of the wild potato and of the potato cultivar ‘Gala’ during these experiments. In downstream feeding experiments, blue stained plant sap was observed in both *S. bulbocastanum* and *S. tuberosum* treated planthopper intestines. This suggests that the planthoppers consumed xylem sap as the plant stalks were cut at the bottom and the top to allow for a quick water transport via transpiration which can be considered to take place in xylem cells (Holbrook and Zwieniecki 2005; Kim et al. 2014). This latter finding adds new insights in the biology of this vector suggesting a flexibility in its feeding strategies which have not been studied so far. At the current status of knowledge, this would be rather non-characteristic for a vector of phloem-limited phytoplasma (Weintraub and Beanland 2006; Gross et al. 2022), but has been observed before in another phloem feeding phytoplasma vector, the psyllid *Cacopsylla picta* (Görg and Gross 2021). Thus, the feeding behavior of *P. leporinus* should be studied in more detail in future feeding experiments.

Most of the planthoppers exposed to *S. bulbocastanum* were dead after a few days of exposure, i.e. about 68% in the first experiment after three days compared to 11% of the control, and about 83% in the second no-choice experiment after two days, compared to 43% of the control. In both experiments, more than 90% of the insects, which fed on *S. bulbocastanum*, were dead after four days. Since the feeding activity was not investigated at individual insect level and in absolute amounts, we can neither conclude nor exclude that this plant has deterrent effects on planthopper feeding behavior. However, choice experiments demonstrated that significantly more insects were observed on the wild potato than on the cultivated potato variety over a two-day period. Other studies would be necessary to investigate which uptake times would be sufficient for toxic effects. Moreover, feeding studies with isolated components such as C16:3-solamine or with plant extracts would be required to infer lethal effects of specific *S. bulbocastanum* compounds on the planthopper. To our knowledge, *P. leporinus* has not been studied yet in terms of behavioral studies, neither exist reports of artificial feeding systems with this species. That would be a pre-requisite for further investigation on toxic effects of acyl solamines in the planthopper.

A proof of the uptake of the plant sap was provided by LC/MS analysis of whole planthoppers and their intestines. We have undertaken this analysis to see whether an ingestion of specific plant ingredients occurred, as those compounds might account for the lower survival rate of the planthoppers on *S. bulbocastanum*. It turned out that C16:3-solamine played a major role in differentiating the groups of planthoppers fed on *S. bulbocastanum* compared to those fed on *S. tuberosum*. In another study, ultrasonication-assisted methanol extraction of dried *S. bulbocastanum* leaves proved to be cytotoxic towards different breast cancer cell lines which was inferred to the phenolic profile of the extract revealing high levels of phenolic acids (Paraschiv et al. 2022). Phenolic acid derivatives were also detected in our LC/MS analysis, but were not causing a significant differentiation of the profiles.

With regards to possible olfactory cues of the wild species leading to attraction of the planthoppers, the VOCs of both species, *S. bulbocastanum* and *S. tuberosum* cv ‘Gala’ have been collected and analyzed. As the planthopper is directed towards the wild crop for at least a first preference, visual or olfactory cues or a combination thereof are likely to play a role in plant detection, as has been reported for other phloem sucking hemipteran insects (Todd et al. 1990; Patt and Sétamou 2007; Wenninger et al. 2009; Aráoz et al. 2019; Gross et al. 2022). In our study, changes in growth stage, plant nutritional status and abiotic factors of the VOC emitting plants were not compared, i.e. it is unclear whether variations of such factors would have resulted in similar results of the VOC profiles or behavioral responses in the choice-experiments. Out of the 80 detected compounds, about half of them (32 + 7) are separating the VOC profiles of the two species. Given this, antennal responses of *P. leporinus* to compounds of these profiles could be further investigated by electroantennography. Subsequent behavioral tests with specific compounds could elicit potential vector behavior to attractant or repellent acting compounds.

Resulting plant infections not only cause damage in yield and quality but also cause concern in potato growing regions, particularly those for seed production and for the food chain. This concern is justified given the reduction of tuber quality in terms of decreased germination capacity and a complete loss of the tradable commodity, but also with regards to critically increased contents of sugars leading to non-processable tubers for the food industry and reduced storage capability (Mahillon et al. 2025; Lang et al. 2025). Consequently, emergency authorizations of insecticides have been approved in Germany in the growing season 2025 for application in potato and sugar beet for the control of *P. leporinus*, but do neither allow for a long-term solution to this pest, nor to the bacterial pathogens transmitted by it.

Given the wide range of host and feeding plants besides potato and sugar beet, this vector poses major challenges for agriculture including the current cropping rotation. Studies of the cropping rotation as a potential to reduce the vector population of *P. leporinus* are sparse (Bressan 2009; Pfitzer et al. 2024). Besides, these studies were conducted before the vector expanded its feeding and host range. Neither the use of repellent crops or substances nor the use of trap crops have been reported so far for *P. leporinus*. Our results suggest that *S. bulbocastanum* is more attractive to *P. leporinus* compared to conventional potato varieties, and at the same time exhibits lethal effects. Given this, *S. bulbocastanum* might serve to attract the vector, and, doing so, reduce it as well as the pathogen population which may decrease because of a lower infection density in field crops. However, this observation lacks proof at field level or at least larger scale, i.e. in field plots. In this scenario, *S. bulbocastanum* might serve as a trap plant, an already existing natural variant of the principle of “attract and kill” (Pickett and Khan 2016; Gregg et al. 2018). With regards to trap plants, Killiny et al. (2021) investigated the effects of biophytoene desaturase-silenced citrus to lure *Diaphorina citri*, the phloem sucking vector of the Huonglongbing disease. Silenced plants expressed visual, olfactory, and gustatory cues and led to a higher attraction of the psyllid vector compared to other citrus plants of the orchard (Killiny et al. 2021). Chaste tree (*Vitex agnus-castus*) was studied as a trap plant in vineyards for *Hyalesthes obsoletus* [Hemiptera: Cixiidae], a planthopper of the same family as *P. leporinus* and also a vector of the phytoplasma ‘*Ca*. Phytoplasma solani’ in grapevine and potato (Maixner 1994; Ulrich et al. 2010). Finally, chaste tree was not recommended as it takes up the transmitted pathogens and acts as a source of infection itself (Moussa et al. 2019). With regards to trap plants and traps in general, there is a knowledge gap for *P. leporinus*. The vector could be attracted at adult stage aboveground but also belowground at nymphal stage according to its lifecycle (Behrmann et al. 2022). With regards to the economic impacts and the need for infection controls, it would be a very interesting question, if *S. bulbocastanum* itself could be infected by ‘*Ca*. Phytoplasma solani’ and ‘*Ca*. Arsenophonus phytopathogenicus’, and if bacteria could proliferate in *S. bulbocastanum* and if symptoms would occur.

Since this planthopper not only expanded its feeding and host plants, but also its spatial distribution, various levers are expected to be used in its future management. Tolerant and resilient crops as well as biotechnological plant protection applications that make use of manipulating vector behavior or interfering vector-plant-interactions would form part of such a strategy. Also, commercially available potato cultivars already possessing genes from *S. bulbocastanum* could be screened for increased pathogen tolerance or effects on the vector.

## Conclusion

To our knowledge, results of *P. leporinus* and its interaction with a wild potato, *S. bulbocastanum*, are presented for the first time by this study. These results could be a starting point for further research directed towards natural plant defense and sustainable crop protection such as potato resistance breeding. *S. bulbocastanum* plants, the application of its leaf extracts or volatile organic compounds could be further investigated for attract-and-kill strategies of the vector *P. leporinus*. Their potential as plants for intercropping between potato ridges or as plants bordering potato fields could be investigated in field trials, too.

## Supporting information

Online Resource 1

## Notes

The authors declare no conflicted or financial interests.

## Data Availability

The datasets generated during and/or analyzed during the current study are available from the corresponding author on reasonable request.

## Acknowledgements

We thank David Löffler and Sanela Kugler for their support in catching planthoppers for the trials, as well as the TH Bingen, Germany for allowing us to collect planthoppers on their fields. Furthermore, we are grateful to Sanela Kugler and Svenja Stein for their excellent assistance with the experiments, Ali Karimi and Alicia Koßmann for their assistance in setting up the GC/MS analysis, and Jannicke Gallinger, Dominik Schmidt and Andreas Jürgens for their support in statistical questions. We thank Roman Gäbelein for the provision of tubers for the experiments. We would like to express our appreciation to Astrid Eben for her input for the plant sap assay.

## Author Contribution Statement

ET, JG and KG conceived and designed research. ET and KG conducted experiments. ET and KG analyzed data. ET wrote the manuscript to which KG contributed sections related to LC/MS analysis. JG, KG and ET revised the manuscript. All authors read and approved the final manuscript.

## References

Agrios GN (2005) Plant Pathology. Elsevier, 5th ed.

Anderson MJ (2001) A new method for non-parametric multivariate analysis of variance. Austral Ecol 26:32–46. 10.1111/j.1442-9993.2001.01070.pp.x

Anderson MJ, Ellingsen KE, McArdle BH (2006) Multivariate dispersion as a measure of beta diversity. Ecol Lett 9:683–693. 10.1111/j.1461-0248.2006.00926.x

Aráoz MVC, Jacobi VG, Fernandez PC, et al (2019) Volatiles mediate host-selection in the corn hoppers Dalbulus maidis (Hemiptera: Cicadellidae) and Peregrinus maidis (Hemiptera: Delphacidae). Bull Entomol Res 109:633–642. 10.1017/S000748531900004X

Bakhsh A, Jabran K, Nazik N, Çalışkan ME (2023) Chapter 25 −Conclusions and future prospective in potato production. In: Çalişkan ME, Bakhsh A, Jabran K (eds) Potato Production Worldwide. Academic Press, pp 457–470

Behrmann SC, Rinklef A, Lang C, et al (2023) Potato (Solanum tuberosum) as a New Host for Pentastiridius leporinus (Hemiptera: Cixiidae) and Candidatus Arsenophonus Phytopathogenicus. Insects 14:281. 10.3390/insects14030281

Behrmann SC, Witczak N, Lang C, et al (2022) Biology and Rearing of an Emerging Sugar Beet Pest: The Planthopper Pentastiridius leporinus. Insects 13:656. 10.3390/insects13070656

Biedermann, Robert, Niedringhaus, Rolf (2009) The Plant- and Leafhoppers of Germany - Identification Key to all Species. Scheeßel: WABV

Bressan A (2009) Agronomic practices as potential sustainable options for the management of Pentastiridius leporinus (Hemiptera: Cixiidae) in sugar beet crops. J Appl Entomol 133:760–766. 10.1111/j.1439-0418.2009.01407.x

Bressan A, Moral García FJ, Sémétey O, Boudon-Padieu E (2010) Spatio-temporal pattern of Pentastiridius leporinus migration in an ephemeral cropping system. Agric For Entomol 12:59–68. 10.1111/j.1461-9563.2009.00450.x

Brooks ME, Kristensen K, Benthem KJvan, et al (2017) glmmTMB Balances Speed and Flexibility Among Packages for Zero-inflated Generalized Linear Mixed Modeling. R J 9:378–400

Brückner A, Heethoff M (2016) A chemo-ecologists’ practical guide to compositional data analysis. 33–46

Crossley MS, Schoville SD, Haagenson DM, Jansky SH (2018) Plant Resistance to Colorado Potato Beetle (Coleoptera: Chrysomelidae) in Diploid F2 Families Derived From Crosses Between Cultivated and Wild Potato. J Econ Entomol 111:1875–1884. 10.1093/jee/toy120

EFSA (2024) Minutes of the 32nd Meeting of the Stakeholder Discussion Group on Emerging Risks

European Commission (2022) Plant pest prevention through technology-guided monitoring and site-specific control | PURPEST | Projekt | Fact Sheet | HORIZON. In: CORDIS Eur. Comm. https://cordis.europa.eu/project/id/101060634. Accessed 3 May 2025

Eurostat (2024) The EU potato sector - statistics on production, prices and trade. https://ec.europa.eu/eurostat/statistics-explained/index.php?title=The_EU_potato_sector_-_statistics_on_production,_prices_and_trade. Accessed 3 May 2025

FAO (2024) Agricultural production statistics 2010–2023

Gallinger J, Jarausch B, Jarausch W, Gross J (2020) Host plant preferences and detection of host plant volatiles of the migrating psyllid species Cacopsylla pruni, the vector of European Stone Fruit Yellows. J Pest Sci 93:461–475. 10.1007/s10340-019-01135-3

Gopal J (2023) Status and way-forward in breeding potato (Solanum tuberosum) for resistance to late blight. Indian J Agric Sci 93:03–10. 10.56093/ijas.v93i1.119721

Görg, LM, Gross J (2021) Influence of ontogenetic and migration stage on feeding behavior of Cacopsylla picta on ‘Candidatus Phytoplasma mali’ infected and non-infected apple plants. J Insect Physiol 131:104229

Gorzolka K, Böttcher C, Thieme T, et al (2025) Characterization and Potential Role of Fatty Acid-Solamine Conjugates in Plant Defense against Herbivores and Pathogens in Wild Potato Species. J Agric Food Chem. 10.1021/acs.jafc.5c01297

Gregg PC, Socorro APD, Landolt PJ (2018) Advances in Attract-and-Kill for Agricultural Pests: Beyond Pheromones. Annu Rev Entomol 63:453–470. 10.1146/annurev-ento-031616-035040

Gross J, Gallinger J, Görg L (2022) Interactions between phloem-restricted bacterial plant pathogens, their vector insects, host plants, and natural enemies, mediated by primary and secondary plant metabolites. Entomol Gen. 10.1127/entomologia/2021/1254

Gross J, Gallinger J, Rid M (2019) Collection, Identification, and Statistical Analysis of Volatile Organic Compound Patterns Emitted by Phytoplasma Infected Plants. In: Musetti R, Pagliari L (eds) Phytoplasmas: Methods and Protocols. Springer, New York, NY, pp 333–343

Guijas C, Montenegro-Burke JR, Domingo-Almenara X, et al (2018) METLIN: A Technology Platform for Identifying Knowns and Unknowns | Analytical Chemistry. Anal Chem 90:3156–3164. 10.1021/acs.analchem.7b04424

Holbrook NM, Zwieniecki MA (2005) Vascular transport in plants. Elsevier Academic Press, Amsterdam

Jansky SH, Spooner DM (2018) The Evolution of Potato Breeding. In: Plant Breeding Reviews, First Edition. John Wiley & Sons, Inc, pp 169–214

Karimi A, Gross J (2024) Development and validation of an innovative headspace collection technique: volatile organic compound patterns emitted by different developmental stages of Halyomorpha halys. Front Hortic 3:. 10.3389/fhort.2024.1380008

Killiny N, Nehela Y, George J, et al (2021) Phytoene desaturase-silenced citrus as a trap crop with multiple cues to attract Diaphorina citri, the vector of Huanglongbing. Plant Sci 308:110930. 10.1016/j.plantsci.2021.110930

Kim HK, Park J, Hwang I (2014) Investigating water transport through the xylem network in vascular plants. J Exp Bot 65:1895–1904. 10.1093/jxb/eru075

Lang C, Dettweiler A, Benaouda S, et al (2025) Pentastiridius leporinus as a plant disease vector: The practical state of knowledge and derived research objectives. Sugar Ind Int 150:105–120. 10.36961/si33023

Lenz M-S, Müller S, Gäbelein R, et al (2025) Bioactivity of acyl solamines from leaves of Solanum bulbocastanum against Alternaria solani and Botrytis cinerea. Potato Res

Lindner K, Haase NU, Roman M, Seemüller E (2011) Impact of Stolbur Phytoplasmas on Potato Tuber Texture and Sugar Content of Selected Potato Cultivars. Potato Res 54:267–282. 10.1007/s11540-011-9192-3

LLH (2024). Landesbetrieb Landwirtschaft Hessen. Angemeldete Vermehrungsflächen Pflanzkartoffel Hessen (ha)

Mahillon M, Bussereau F, Dubuis N, et al (2025) First Detection of Arsenophonus in Potato Crop in Switzerland: A Threat for the Processing Industry? Potato Res. 10.1007/s11540-024-09840-y

Maixner M (1994) Transmission of German grapevine yellows (Vergilbungskrankheit) by the planthopper Hyalesthes obsoletus (Auchenorrhyncha: Cixiidae). VITIS - J Grapevine Res 33:103–103. 10.5073/vitis.1994.33.103-104

Makowski D, Ben-Shachar MS, Wiernik BM, et al (2025) modelbased: An R package to make the most out of your statistical models through marginal means, marginal effects, and model predictions. J Open Source Softw 10:7969. 10.21105/joss.07969

Malcolmson JF, Black W (1966) New R genes in Solanum demissum lindl. And their complementary races of Phytophthora infestans (Mont.) de bary. Euphytica 15:199–203. 10.1007/BF00022324

McGillycuddy M, Popovic G, Bolker BM, Warton DI (2025) Parsimoniously Fitting Large Multivariate Random Effects in glmmTMB. J Stat Softw 112:1–19. 10.18637/jss.v112.i01

Moussa A, Mori N, Faccincani M, et al (2019) Vitex agnus-castus cannot be used as trap plant for the vector Hyalesthes obsoletus to prevent infections by ‘Candidatus Phytoplasma solani’ in northern Italian vineyards: Experimental evidence. Ann Appl Biol 175:302–312. 10.1111/aab.12542

NIST Chemistry WebBook (2025), NIST Standard Reference Database Number 69, National Institute of Standards and Technology (NIST), United States of America. 10.18434/T4D303

Oksanen J, Simpson GL, Blanchet FG, et al (2025) vegan: Community Ecology Package

Pang Z, Chong J, Zhou G, et al (2021) MetaboAnalyst 5.0: narrowing the gap between raw spectra and functional insights. Nucleic Acids Res 49:W388–W396. 10.1093/nar/gkab382

Pang Z, Zhou G, Ewald J, et al (2022) Using MetaboAnalyst 5.0 for LC–HRMS spectra processing, multi-omics integration and covariate adjustment of global metabolomics data. Nat Protoc 17:1735–1761. 10.1038/s41596-022-00710-w

Paraschiv M, Csiki M, Diaconeasa Z, et al (2022) Phytochemical Profile and Selective Cytotoxic Activity of a Solanum bulbocastanum Dun. Methanolic Extract on Breast Cancer Cells. Plants 11:3262. 10.3390/plants11233262

Patt JM, Sétamou M (2007) Olfactory and Visual Stimuli Affecting Host Plant Detection in Homalodisca coagulata (Hemiptera: Cicadellidae). Environ Entomol 36:142–150. 10.1603/0046-225X(2007)36[142:OAVSAH]2.0.CO;2

Pfitzer R, Rostás M, Häußermann P, et al (2024) Effects of succession crops and soil tillage on suppressing the syndrome ‘basses richesses’ vector Pentastiridius leporinus in sugar beet. Pest Manag Sci 80:3379– 3388. 10.1002/ps.8041

Pickett JA, Khan ZR (2016) Plant volatile-mediated signalling and its application in agriculture: successes and challenges. New Phytol 212:856–870. 10.1111/nph.14274

Rid M, Mesca C, Ayasse M, Gross J (2016) Apple Proliferation Phytoplasma Influences the Pattern of Plant Volatiles Emitted Depending on Pathogen Virulence. Front Ecol Evol 3:. 10.3389/fevo.2015.00152

Rinklef A, Behrmann SC, Löffler D, et al (2024) Prevalence in Potato of ‘Candidatus Arsenophonus Phytopathogenicus’ and ‘Candidatus Phytoplasma Solani’ and Their Transmission via Adult Pentastiridius leporinus. Insects 15:275. 10.3390/insects15040275

Tais L, Schulz H, Böttcher C (2021) Comprehensive profiling of semi□polar phytochemicals in whole wheat grains (Triticum aestivum) using liquid chromatography coupled with electrospray ionization quadrupole time□of□flight mass spectrometry. Metabolomics 17:18. 10.1007/s11306-020-01761-4

Takeshita K, Kikuchi Y (2020) Genomic Comparison of Insect Gut Symbionts from Divergent Burkholderia Subclades. Genes 11:744. 10.3390/genes11070744

Therhaag E, Schneider B, Zikeli K, et al (2024a) Pentastiridius leporinus (Linnaeus, 1761) as a Vector of Phloem-Restricted Pathogens on Potatoes: ‘Candidatus Arsenophonus Phytopathogenicus’ and ‘Candidatus Phytoplasma Solani.’ Insects 15:189. 10.3390/insects15030189

Therhaag E, Ulrich R, Gross J, Schneider B (2024b) Onion (Allium cepa) as a New Host for ‘Candidatus Arsenophonus phytopathogenicus’ in Germany. Plant Dis 108:2914. 10.1094/PDIS-03-24-0526-PDN

Therneau TM, until 2009) TL (original S->R port and R maintainer, Elizabeth A, Cynthia C (2024) survival: Survival Analysis

Todd JL, Phelan PL, Nault LR (1990) Interaction between visual and olfactory stimuli during host-finding by leafhopper,Dalbulus maidis (Homoptera: Cicadellidae). J Chem Ecol 16:2121–2133. 10.1007/BF01026924

Ulrich R, Preiss U, Fabich S (2010) Potato Stolbur phytoplasma in Hesse and Rhineland-Palatinate. Julius-Kühn-Arch

Van Der Vossen E, Sikkema A, Hekkert B te L, et al (2003) An ancient R gene from the wild potato species Solanum bulbocastanum confers broad-spectrum resistance to Phytophthora infestans in cultivated potato and tomato. Plant J 36:867–882. 10.1046/j.1365-313X.2003.01934.x

Weintraub PG, Beanland L (2006) INSECT VECTORS OF PHYTOPLASMAS. Annu Rev Entomol 51:91–111. 10.1146/annurev.ento.51.110104.151039

Wenninger EJ, Stelinski LL, Hall DG (2009) Roles of Olfactory Cues, Visual Cues, and Mating Status in Orientation of Diaphorina citri Kuwayama (Hemiptera: Psyllidae) to Four Different Host Plants. Environ Entomol 38:225–234. 10.1603/022.038.0128

Wickham H (2016) ggplot2: Elegant Graphics for Data Analysis. Springer-Verlag New York

Xia J, Psychogios N, Young N, Wishart DS (2009) MetaboAnalyst: a web server for metabolomic data analysis and interpretation. Nucleic Acids Res 37:W652–W660. 10.1093/nar/gkp356

Zaheer K, and Akhtar MH (2016) Potato Production, Usage, and Nutrition—A Review. Crit Rev Food Sci Nutr 56:711–721. 10.1080/10408398.2012.724479

