## Supplementary material for "“Lethal effects of the wild potato *Solanum bulbocastanum* on the planthopper *Pentastiridius leporinus*, a vector of bacterial pathogens in potato”": Online Resource 1

**Online Resource 1** Table of numbered, identified, annotated or unknown volatile organic compounds (C1 - C80) found in *S. bulbocastanum* or *S. tuberosum* cv 'Gala' or in both species, indicating the retention indices (RI)

| $C_i$ | Compound | RI | Species |
|---|---|---|---|
| C1 | Hexanal [b] | 798 | both |
| C2 | trans-2-Hexenal [b] | 849 | " |
| C3 | 3-Hexen-1-ol, (E)- [a] | 852 | " |
| C4 | 1-Nonene [a] | 890 | " |
| C5 | α-Pinene [b] | 932 | *S. tuberosum cv 'Gala'* |
| C6 | Benzaldehyd [b] | 957 | both |
| C7 | Benzonitrile [a] | 982 | " |
| C8 | 5-Hepten-2-one, 6-methyl- [b] | 987 | " |
| C9 | β –Myrcene [b] | 991 | *S. tuberosum cv 'Gala'* |
| C10 | Decane [b] | 1000 | both |
| C11 | Octanal [b] | 1003 | " |
| C12 | 3-Hexen-1-ol, acetate, (Z)- [b] | 1007 | " |
| C13 | α-Terpinene [a] | 1016 | *S. tuberosum cv 'Gala'* |
| C14 | (R)-(+)-Limonene [b] | 1028 | *S. tuberosum cv 'Gala'* |
| C15 | unknown_RI_1043 (*m/z* 87, 102, 57, 41, 130) | 1043 | both |
| C16 | Ocimene [b] | 1048 | " |
| C17 | unknown_RI_1067 (*m/z* 57, 41, 43, 56) | 1067 | " |
| C18 | Methylbenzoat [b] | 1094 | " |
| C19 | Undecane [b] | 1100 | " |
| C20 | Pelargonaldehyd [b] | 1104 | " |
| C21 | 2-Phenylethanol [b] | 1112 | *S. bulbocastanum* |
| C22 | 4,8-dimethyl-1(E)3,7-nonatriene [b] | 1117 | *S. tuberosum cv 'Gala'* |
| C23 | unknown_RI_1159 (*m/z* 73, 267, 268, 45) | 1159 | both |
| C24 | unknown_RI_1169 (*m/z* 57, 43, 41, 82, 69) | 1169 | " |
| C25 | 1-Dodecene [a] | 1192 | " |
| C26 | Methyl salicylate [b] | 1194 | " |
| C27 | Decanal [b] | 1206 | " |
| C28 | Benzothiazol [b] | 1222 | *S. tuberosum cv 'Gala'* |
| C29 | Nonanoic acid [a] | 1275 | both |
| C30 | Undecanal [b] | 1307 | " |

**Online Resource 1** *continued*

| $C_i$ | Compound | RI | Species |
|---|---|---|---|
| C31 | unknown_RI_1326 (*m/z* 71, 43, 57, 85, 41) | 1326 | both |
| C32 | unknown_RI_1343 (*m/z* 83, 41, 57, 95, 109) | 1343 | “ |
| C33 | unknown_RI_1339 (*m/z* 121, 93, 107, 79, 41) | 1339 | *S. tuberosum cv ‘Gala’* |
| C34 | α-Cubebene [a] | 1351 | *S. tuberosum cv ‘Gala’* |
| C35 | unknown_RI_1363 (*m/z* 43, 57, 41, 70, 71) | 1363 | *S. bulbocastanum* |
| C36 | (-)-α-Copaene [a] | 1378 | *S. tuberosum cv ‘Gala’* |
| C37 | (-)-β-Bourbonene [a] | 1387 | *S. tuberosum cv ‘Gala’* |
| C38 | 1-Tetradecene [a] | 1392 | *S. bulbocastanum* |
| C39 | Cyclohexane [a] | 1394 | both |
| C40 | Tetradecane [b] | 1399 | “ |
| C41 | Dodecanal [a] | 1409 | “ |
| C42 | α-Gurjunene [a] | 1413 | *S. tuberosum cv ‘Gala’* |
| C43 | β-Caryophyllene [b] | 1424 | *S. tuberosum cv ‘Gala’* |
| C44 | trans-α-Bergamotene [a] | 1438 | *S. tuberosum cv ‘Gala’* |
| C45 | unknown_RI_1445 (*m/z* 69, 41, 91, 55, 77) | 1445 | *S. tuberosum cv ‘Gala’* |
| C46 | Geranyl acetone [b] | 1453 | *S. tuberosum cv ‘Gala’* |
| C47 | unknown_RI_1458 (*m/z* 165, 193, 151, 95, 208) | 1458 | *S. bulbocastanum* |
| C48 | (E)-β-Farnesene [a] | 1460 | *S. tuberosum cv ‘Gala’* |
| C49 | cis-Muurola-4(15),5-diene [a] | 1467 | *S. tuberosum cv ‘Gala’* |
| C50 | unknown_RI_1468 (*m/z* 177, 41, 220, 135, 57) | 1468 | both |
| C51 | γ-Muurolene [a] | 1480 | *S. tuberosum cv ‘Gala’* |
| C52 | (-)-Germacrene D [a] | 1486 | *S. tuberosum cv ‘Gala’* |
| C53 | unknown_RI_1496 (*m/z* 71, 57, 43, 85, 41) | 1496 | *S. bulbocastanum* |
| C54 | α-Farnesene [a] | 1510 | *S. tuberosum cv ‘Gala’* |
| C55 | γ-Cadinene [a] | 1518 | *S. tuberosum cv ‘Gala’* |
| C56 | δ-Cadinene [a] | 1527 | *S. tuberosum cv ‘Gala’* |
| C57 | unknown_RI_1541 (*m/z* 71, 57, 43, 85, 41) | 1541 | both |
| C58 | unknown_RI_1543 (*m/z* 55, 54, 71, 43, 41) | 1543 | *S. bulbocastanum* |
| C59 | α-Calacorene [a] | 1547 | *S. tuberosum cv ‘Gala’* |
| C60 | unknown_RI_1572 (*m/z* 105, 67, 111, 122, 41) | 1572 | *S. tuberosum cv ‘Gala’* |
| C61 | (3E,7E)-4,8,12-Trimethyltrideca-1,3,7,11-tetraene [a] | 1580 | both |
| C62 | Caryophyllene oxide [a] | 1588 | *S. tuberosum cv ‘Gala’* |
| C63 | Cetene [a] | 1592 | both |
| C64 | Hexadecane [b] | 1599 | “ |
| C65 | unknown_RI_1608 (*m/z* 41, 67, 82, 111, 71) | 1608 | *S. tuberosum cv ‘Gala’* |
| C66 | α-Corocalene [a] | 1626 | *S. tuberosum cv ‘Gala’* |
| C67 | unknown_RI_1628 (*m/z* 57, 41, 138, 181, 69) | 1628 | both |
| C68 | τ-Cadinol [a] | 1645 | *S. tuberosum cv ‘Gala’* |
| C69 | unknown_RI_1663 (*m/z* 57, 43, 41, 69, 44) | 1663 | both |
| C70 | unknown_RI_1668 (*m/z* 57, 41, 95, 138, 67) | 1668 | “ |
| C71 | unknown_RI_1679 (*m/z* 183, 198, 168, 153, 165) | 1679 | *S. tuberosum cv ‘Gala’* |
| C72 | unknown_RI_1698 (*m/z* 57, 43, 71, 85) | 1698 | both |
| C73 | unknown_RI_1703 (*m/z* 207, 235, 57, 250, 222) | 1703 | “ |
| C74 | unknown_RI_1710 (*m/z* 71, 57, 43, 85, 69) | 1710 | “ |
| C75 | unknown_RI_1734 (*m/z* 43, 91, 79, 121, 55) | 1734 | *S. tuberosum cv ‘Gala’* |
| C76 | unknown_RI_1756 (*m/z* 71, 57, 43, 85, 99) | 1756 | both |
| C77 | unknown_RI_1771 (*m/z* 219, 191, 234, 57, 220) | 1771 | “ |

**Online Resource 1** *continued*

| $C_i$ | Compound | RI | Species |
|---|---|---|---|
| C78 | unknown_RI_1793 (*m/z* 55, 83, 97, 41, 57) | 1793 | both |
| C79 | unknown_RI_1844 (*m/z* 69, 41, 81, 55, 93) | 1844 | *S. tuberosum cv 'Gala'* |
| C80 | unknown_RI_1972 (*m/z* 69, 93, 41, 81, 107) | 1972 | *S. tuberosum cv 'Gala'* |

[a] Annotated compounds derived from NIST Mass Spectral Search Program and comparison of retention index (RI) data from NIST Chemical WebBook

[b] Compounds identified by comparing mass spectrum and RI with authentic standards on the same GC-MS

Mass fragments for unknowns are listed with the base peak and other fragments in order of decreasing abundance
